# Vitamin D status is heritable and under environment-dependent selection in the wild

**DOI:** 10.1101/2020.08.14.251108

**Authors:** Alexandra M. Sparks, Susan E. Johnston, Ian Handel, Jill G. Pilkington, Jacqueline Berry, Josephine M. Pemberton, Daniel H. Nussey, Richard J. Mellanby

## Abstract

Vitamin D has a well-established role in skeletal health and is increasingly linked to chronic disease and mortality in humans and companion animals. Despite the clear significance of vitamin D for health and obvious implications for fitness under natural conditions, no longitudinal study has tested whether the circulating concentration of vitamin D is under natural selection in the wild. Here, we show that concentrations of dietary-derived vitamin D_2_ and endogenously-produced vitamin D_3_ metabolites are heritable and largely polygenic in a wild population of Soay sheep (*Ovis aries*). Vitamin D_2_ status was positively associated with female adult survival, and vitamin D_3_ status predicted female fecundity in particular, good environment years. Our study provides evidence that vitamin D status has the potential to respond to selection, as well as new insights into how vitamin D metabolism is associated with fitness in the wild.

## Introduction

Vitamin D is a critical component in the development and maintenance of skeletal health ^1^. However, current research suggests that vitamin D may also have wider biomedical effects, with studies linking vitamin D insufficiency to increased risk of mortality in humans and companion animals ^2–4^, as well as reproductive failure, low birth weight, infertility and reduced litter sizes in humans, rats and mice ^5–8^. This link suggests that vitamin D status may be associated with fitness in natural populations, yet there have been no investigations into the causes and consequences of its variation in wild populations. To understand the ecological and evolutionary significance of vitamin D metabolism, it is necessary to determine how it varies due to individual and environmental variation, the extent to which it is heritable and the underlying genetic architecture, and its association with key components of fitness, such as survival and fecundity. By investigating these factors, we can begin to understand the evolutionary potential of vitamin D status and determine if it is under selection in natural conditions.

The availability of vitamin D to mammals is dependent on the environment. Vitamin D_3_ can be produced in the skin of most mammals following exposure to sunlight. Vitamin D_3_ can also be obtained from dietary sources, notably oily fish, eggs and liver, whereas Vitamin D_2_ can only be obtained in the diet from some plants and fungi ^1^. Human studies suggest vitamin D status, typically assessed by measuring circulating concentrations of the major metabolite 25 hydroxyvitamin D (25(OH)D) is modestly heritable and genome-wide association studies have identified several genes involved in relevant pathways which contribute significantly to this genetic variation ^9–16^. However, vitamin D supplementation and artificial exposure to ultraviolet light are increasingly common in Western human populations, whilst variation in challenging aspects of the environment (e.g. thermoregulation, infection, food limitation) are greatly reduced compared to most natural populations ^17^. As a result, the potential for genes identified in human studies to contribute to variation in vitamin D metabolism and thus potentially be under natural selection more broadly, remains unknown.

Here, we investigate the genetic architecture and fitness consequences of vitamin D status in a long-term study of wild Soay sheep on the St Kilda archipelago in NW Scotland. A previous small-scale study of females sampled in a single year showed that plasma 25(OH)D_2_ and 25(OH)D_3_ concentrations varied among individuals, and identified associations between vitamin D status, age, coat colour and female fecundity ^18^. In the present study, we use samples collected longitudinally over a six-year period, alongside detailed information on the genetics and life histories of the individuals sampled, to: (i) estimate the heritability of vitamin D status in the wild; (ii) use a genome-wide association study (GWAS) approach to determine the genetic architecture of vitamin D status, and (iii) examine how vitamin D status predicts survival and reproductive performance.

## Methods

### Study population

The Soay sheep is a primitive breed of domestic sheep that was isolated on the island of Soay in the remote St Kilda archipelago (57°49′N 8°35′W) several millennia ago, and has been living wild since then ^19^. In 1932, just over 100 Soay sheep were moved to the larger island of Hirta after the evacuation of all human residents and livestock. The population expanded and now fluctuates between 600 and 2,200 sheep. Approximately one third of the Hirta population lives in the Village Bay area, and these individuals have been the subject of a long-term study since 1985 ^19^. In April, around 95% of all lambs born in this area are caught each year and individually tagged; most females have a single lamb per year, with twins occurring in ∼13% of births. Each August, as many sheep as possible from the study population are re-captured using temporary traps ^19^. At capture, animals are weighed, and blood samples are collected. Whole blood samples are collected into heparin tubes, centrifuged at 3000 r.p.m. for 10 minutes, and plasma removed and stored at −20°C. This analysis included all animals that were caught and blood sampled in August between 2011 and 2016, comprising 1452 samples from 917 individuals. A breakdown of the dataset by sex, age group and year is shown in Table S1.

### Vitamin D measurements

In the Soay sheep population on Hirta, Vitamin D_2_ is available from dietary sources and vitamin D_3_ is only obtained from cutaneous production ^19,20^. This population of Soay sheep do not receive any diet supplementation, therefore we are confident that vitamin D_2_ was obtained via grazing, whereas vitamin D_3_ was only obtained from cutaneous production. Vitamin D is initially metabolised to 25(OH)D in the liver before undergoing a second hydroxylation in the kidney to the metabolically active 1,25 dihydroxyvitamin D (1,25(OH)_2_D). Circulating concentrations of 25(OH)D are widely used to assess vitamin D status in mammals ^1^. Plasma concentrations of 25(OH)D_2_ and 25(OH)D_3_ were determined by liquid chromatography tandem mass spectrophotometry (LC-MS/MS) using an ABSciex 5500 tandem mass spectrophotometer (Warrington, UK) and the Chromsystems (Munich, Germany) 25OHD kit for LC-MS/MS following the manufacturers’ instructions (intra- and inter-assay CV 3.7% and 4.8% respectively) in a laboratory certified as proficient by the international Vitamin D Quality Assurance Scheme (DEQAS). Total 25(OH)D was defined as the sum of 25(OH)D_2_ and 25(OH)D_3_.

### Phenotypic associations with vitamin D concentrations

We first investigated phenotypic associations with total 25(OH)D, 25(OH)D_2_ and 25(OH)D_3_ concentrations using a linear mixed-effects model in the lme4 package v1.1-21 ^21^ in R v3.5.3. We included sex (factor), age (years, linear and quadratic), coat colour (factor: light/dark), and year (factor) as fixed effects and individual identity and birth year as random effects. Significance of fixed effects were determined by dropping each fixed effect from a model containing all terms and performing a likelihood ratio test.

### SNP and pedigree data set

A total of 7,700 Soay sheep have been genotyped at 51,135 SNP markers on the Illumina Ovine SNP50 BeadChip. Quality control was carried out using the *check*.*marker* function in GenABEL v1.8-0 ^22^ in R v3.4.3 using the following thresholds: SNP minor allele frequency (MAF) > 0.001, SNP locus genotyping success > 0.99, individual sheep genotyping success > 0.95; identity by state with another individual < 0.9. Following quality control, 39,398 SNPs remained. All SNP locations were taken from their estimated positions on the sheep genome assembly Oar_v3.1 (GenBank assembly ID GCA_000298735.1) ^23^. A pedigree has previously been inferred using 438 unlinked SNPs in the R package Sequoia ^24^; in cases where SNP information was not available, then parentage was inferred from field observations (for mothers) or from microsatellite data ^25^. We genotyped 189 sheep at 606,066 SNP loci on the Illumina Ovine Infinium HD SNP BeadChip; these sheep were selected to maximise the genetic diversity represented in the full population to allow for genotype imputation (see ^26^ for full details). A total of 430,702 SNPs for 188 individuals passed quality control using the thresholds above. Briefly, SNP genotypes from the HD chip were imputed into the 50K typed individuals using AlphaImpute v1.98 ^27,28^, resulting in a dataset with 7691 individuals, 420,556 SNPs and a mean genotyping rate per individual of 99.5% (range 94.8% - 100%; see ^29^ for full details). All genotype positions from both SNP chips were based on the Oar_v3.1 sheep genome assembly. A total of 900 out of 917 unique individuals with vitamin D measures had corresponding genomic data.

### Animal models

We fitted quantitative genetic “animal models” in order to determine the heritability and repeatability of total 25(OH)D, 25(OH)D_2_ and 25(OH)D_3_ concentrations in ASReml-R 4.1.0 ^30^ in R v3.6.2. Using the above SNP dataset, a genomic relatedness matrix (GRM) at all autosomal markers on the SNP50 Chip was constructed for all 7,700 genotyped individuals using GCTA v1.91.1 ^31^ with the argument *-- grm-adj 0* which assumes that causal and genotyped loci have similar frequency spectra. Univariate animal models were fitted for each of the three vitamin D measures across all ages. The fixed effect structure for the model included sex and year of capture as factors, and age as a linear and quadratic covariate. The random effects included the additive genetic component (using the GRM), permanent environment component (due to repeated measures within an individual), maternal identity and birth year. Significance of random effects were determined by dropping each random effect from a model containing all random effects and performing a likelihood ratio test distributed as. The proportion of the phenotypic variance explained by each random effect was estimated as the ratio of the relevant variance component to total phenotypic variance. The heritability of each measure was determined as the ratio of the additive genetic variance to the total phenotypic variance. The repeatability (i.e. the between-individual variation) of each measure was determined as the ratio of the sum of the additive genetic and permanent environment variance to the total phenotypic variance (equivalent to the proportion of variance explained by individual identity).

Bivariate animal models were fitted with 25(OH)D_2_ and 25(OH)D_3_ concentrations as the two response variables to estimate covariances at the additive genetic (using the GRM), permanent environment and residual levels. Fixed effects were fitted as above. Covariance components were summed to estimate the total phenotypic covariance between each pair of traits. Correlations were determined by dividing the covariance by the product of the square roots of the two variances at the additive genetic, permanent environment and residual level. Significance of the genetic correlation was determined by running the bivariate model and using the *corgh* function to fix the genetic correlation at 0 or 1 (0.999) and comparing with the observed model using a likelihood ratio test.

### Genome-wide association study

A genome-wide association study using the imputed SNP dataset was used to determine associations between individual SNPs and the three vitamin D concentrations using the R package RepeatABEL v1.1 ^32^ in R v 3.4.3. First, a linear mixed-effect model was constructed using the same model structures as for the animal models above using the function *preFitModel*. Then, the function rGLS was used to fit SNP effects in the model. This approach fits both repeated measures and the GRM, which will account for pseudoreplication and population structure, respectively ^32^. The association P-values were corrected for any additional unaccounted population structure by dividing them by the genomic control parameter λ, in cases where λ > 1, to reduce the incidence of false positives. The threshold for genome-wide significance was previously determined using linkage disequilibrium information to determine the effective number of independent tests ^29^; the genome-wide significance threshold was set at P < 1.28 × 10^−6^ at α = 0.05. In genome regions where a SNP locus was significantly associated with vitamin D, we first estimated the genotype effect sizes by fitting the genotype as a three-level factor in the animal model structures above. Then, in separate models, the proportion of phenotypic variance explained by that region was modelled as follows: a second “regional” GRM was constructed using 10 flanking 50K-SNP loci on either side (i.e. at least 20 SNPs in total) or the first/last 20 50K-SNPs of the chromosome if the number of flanking SNPs on one side was <10; this regional GRM was then fit as an additional random effect in the animal model above and significance determined using a likelihood ratio test as above.

In order to identify potential candidate genes for vitamin D concentrations in significant regions, sheep gene IDs and their associated gene ontology (GO annotations) within 1Mb of the most highly associated SNPs (P < 10^−5^) were extracted from Ensembl (gene build ID Oar_v3.1.100) using the function *getBM* in the R package biomaRt v2.34.2 ^33^. Gene orthologues in humans (*Homo sapiens*), cattle (*Bos taurus*), mouse (*Mus musculus*) and rat (*Rattus norvegicus*) and associated GO terms were also extracted using the function *getLDS*. Gene and orthologue names, descriptions, phenotype descriptions and GO terms were then queried for terms associated with 25(OH)D concentrations in human studies ^11^, using the R command *grep* with the strings *vitamin D, cholecalcif* (Vitamin D_2_), *ergoecalcif* (Vitamin D_3_), *CYP2, NADSYN1, DHCR7, SEC23A* and *AMDHD1*.

### Fitness analyses

We investigated associations between total 25(OH)D, 25(OH)D_2_ and 25(OH)D_3_ concentrations measured in summer and survival over the subsequent winter, since the vast majority of sheep deaths occur over the winter months ^19^. Accurate death date information is known from regular censuses and searches for carcasses over this period. Over-winter survival was defined as survival from capture and sampling in August to 1^st^ May in the subsequent year. Due to known differences in survival rates between lambs and adult Soay sheep, we modelled lambs and adults separately (where the adult group included all sheep aged ≥ 1 year). Both lamb and adult survival analyses were conducted using generalised linear models (GLMs) with a binomial error distribution and a logit link function. The lamb base model included sex (factor), coat colour (factor: light/dark), weight (covariate) and year (factor) as fixed effects. The base model for adults included the fixed effects of sex (factor), age group (factor with 3 levels: yearlings, 2-6 years and ≥7 years), coat colour (factor), weight (covariate) and year (factor) and a sex by age group interaction. Covariates were rescaled to mean 0 and standard deviation 1 prior to inclusion in each model. To investigate associations between vitamin D status and over-winter survival, we compared the AIC of this base model with models containing 25(OH)D, 25(OH)D_2_ and 25(OH)D_3_ separately and a model with both 25(OH)D_2_ and 25(OH)D_3_ fitted additively. We also ran these models with interactions between each vitamin D measure and sex, year and age group (adult models only) to test for environment, sex and age dependent associations. The model with the lowest AIC value was considered the best-fitting model, unless the difference in AIC was within 2 AIC units of a model with fewer explanatory terms, in which case this simpler model was considered the most parsimonious ^34^.

Next, we tested for associations between total 25(OH)D, 25(OH)D_2_ and 25(OH)D_3_ concentrations and breeding success in the following spring. For females who survived the winter, annual breeding success was defined as the number of offspring born to the female in the following spring derived from observational data. For males, annual breeding success was calculated as the number of offspring sired in the following spring using the genetic pedigree. Male annual breeding success included all males who died over the winter but excluded all males that were not observed to take part in the rut using census records taken between October-December. Since the distribution of annual breeding success is very different between males and females, and between lambs and adults, models were run for female lambs, male lambs, adult females and adult males separately (where the adult group included all sheep aged ≥ 1 year). For female lambs, breeding success was treated as a binary measure (produced a lamb / did not produce a lamb) as no females had twins in their first year and was run as a GLM with binomial error distribution and logit link function. In this model, fixed effects included coat colour (factor), weight (covariate) and year (factor). Data from 2011 were excluded due to the small number of available measures (n=4) driven by very high lamb over-winter mortality, but the results were consistent with or without these measures (Table S2). For adult females, which have 0-2 lambs per year, annual breeding success analyses were conducted using a proportional odds mixed model in the ordinal package v2019.12-10 using the adaptive Gauss-Hermite quadrature approximation ^35^. Fixed effects included age group (factor, levels as before), coat colour (factor), weight (covariate) and year (factor) with individual identity included as a random effect. Since the probability of males siring offspring in their first year is low (10% in this dataset), male lamb breeding success was treated as a binary measure (sired a lamb/did not sire a lamb) and was run as a GLM with binomial error distribution and logit link function with fixed effects of coat colour (factor), weight and year (factor) as fixed effects. No males born in 2011 in this dataset sired offspring in their first year so this year was excluded from analyses to improve model convergence, although results were consistent with or without these measures (Table S2). Finally, for adult males we analysed annual breeding success in the glmmTMB package v1.0.1 ^36^ where the best-fitting model was the negative binomial model with the “nbinom1” parametrisation without zero inflation. Fixed effects in this model included age group (factor), coat colour (factor), weight and year (factor) with individual identity as a random effect. Continuous variables were rescaled to mean 0 and standard deviation 1 prior to inclusion in each model. As for the survival analyses, AIC of base models were compared with models with 25(OH)D, 25(OH)D_2_, 25(OH)D_3_ separately and 25(OH)D_2_ and 25(OH)D_3_ combined and models where there was a vitamin D measure by year or age group (adult models only) interaction.

### Data availability

Data will be archived on a publicly accessible repository. All scripts for the analysis are provided at https://github.com/sejlab/Soay_Vitamin_D.

## Results

### Individual variation in vitamin D concentrations

25(OH)D_2_ and 25(OH)D_3_ concentrations were positively correlated with one another (Pearson’s product-moment correlation: r=0.59, t_1450_=28.03, p<0.001; Figure S1) but 25(OH)D_3_ concentrations contributed more to the total 25(OH)D concentrations and were on average 26.13 nmol/l higher (Paired t-test: 95% CI 25.04-27.23 nmol/l) than 25(OH)D_2_ concentrations. There was a quadratic association with all vitamin D measures and age (Tables S3 and S4, Figures S2 and S3). In models where age was fitted as a factor, there was a substantial increase in concentrations of the vitamin D metabolites between lambs (Age 0) and yearlings (Age 1) of 35.51 nmol/l (± 1.50 SE), 5.99 nmol/l (± 0.40 SE) and 29.53 nmol/l (± 1.30 SE) for total 25(OH)D, 25(OH)D_2_ and 25(OH)D_3_ respectively. For 25(OH)D_2_, this was followed by a decline with age that was not as apparent for total 25(OH)D and 25(OH)D_3_ (Tables S3 and S4, Figure S2). There was no difference in total 25(OH)D, 25(OH)D_2_ or 25(OH)D_3_ concentrations between the sexes (Table S3). Sheep with light coats had significantly higher total 25(OH)D and 25(OH)D_3_ concentrations of 5.51 nmol/l (±1.35 SE) and 6.25 nmol/l (±1.16 SE) respectively compared to sheep with dark coats (Table S3). Light coated sheep had slightly lower 25(OH)D_2_ concentrations at −0.74 nmol/l (±0.35 SE) than dark coated sheep although this association was only marginally significant (Table S3, p=0.036). All vitamin D measures varied significantly by year (Tables S3 and S4, Figure S3).

There was considerable among-individual variation in each of the three vitamin D measures with repeatabilities of 0.19 for total 25(OH)D, 0.29 for 25(OH)D_2_ and 0.28 for 25(OH)D_3_ (Table S5, Figure 1), where most of this variation (59-87%) was explained by the additive genetic component (see next section). The consistency of vitamin D concentrations within individuals is further illustrated by the strong positive correlation between measures taken from the same individual across consecutive years in adults (Figure S4; Pearson’s product-moment correlations: total 25(OH)D: r = 0.62, t_350_ = 14.89, p < 0.001; 25(OH)D_2_: r = 0.76, t_350_ = 21.82, p < 0.001; 25(OH)D_3_: r = 0.58, t_350_ = 13.21, p < 0.001).

**Figure 1.**
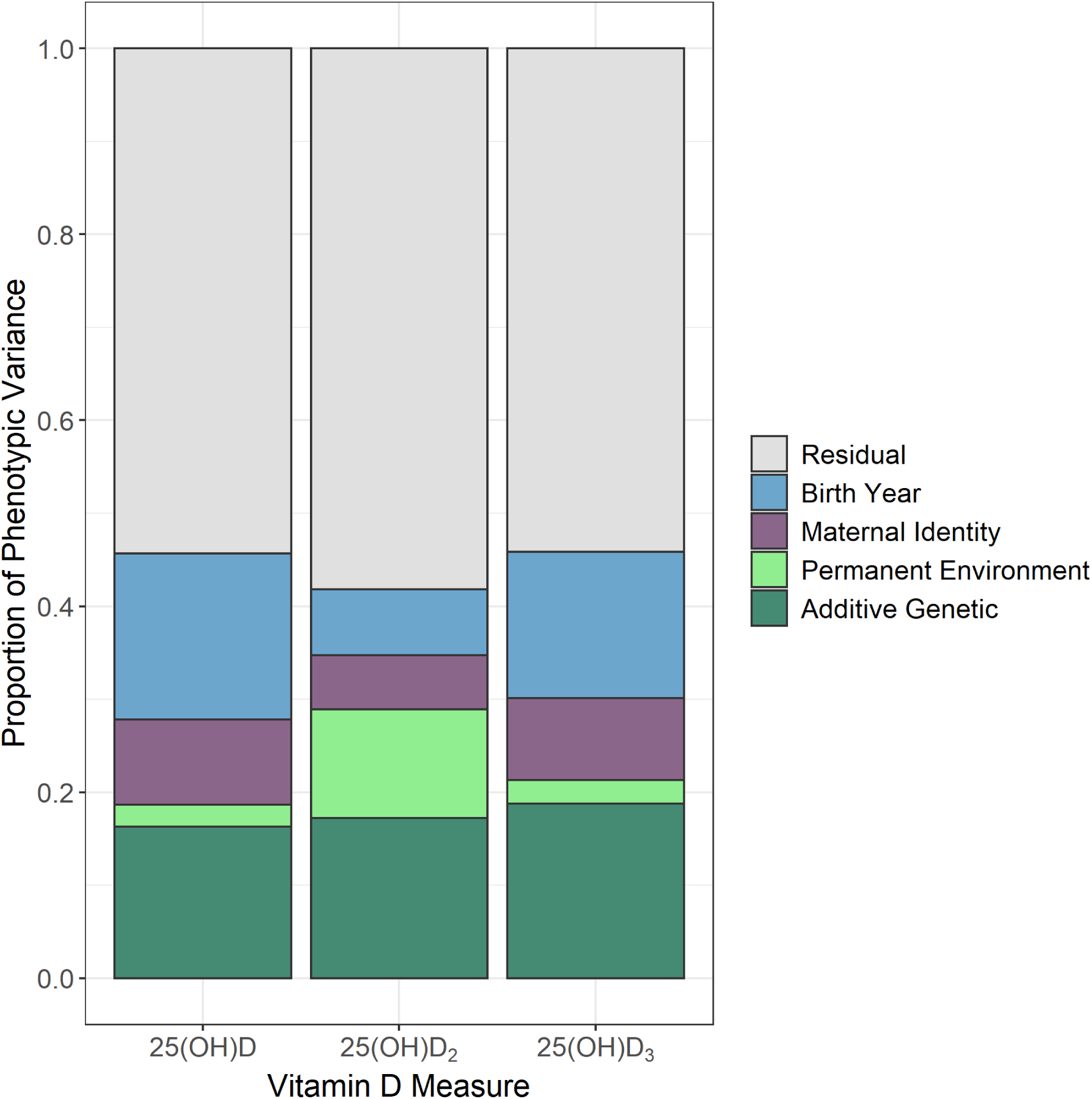
The proportion of phenotypic variance explained by different random effects in univariate animal models for total 25(OH)D, 25(OH)D_2_ and 25(OH)D_3_ plasma concentrations in wild Soay sheep.

### Genetic architecture of vitamin D concentrations

All three vitamin D measures were heritable: h^2^ = 0.16 (± 0.04 SE) for 25(OH)D, h^2^ = 0.17 (± 0.04 SE) for total 25(OH)D_2_ and h^2^ = 0.19 (± 0.04 SE) for 25(OH)D_3_ (Table S5, Figure 1). Variance attributable to birth year ranged from 7.1 – 17.8% across vitamin D measures and there was evidence for maternal effects for total 25(OH)D and 25(OH)D_3_ measures (2.5 – 9.1% of total phenotypic variance; see Figure 1, Table S5). A bivariate model of 25(OH)D_2_ and 25(OH)D_3_ showed there was a moderate positive genetic correlation between the two traits (r_a_=0.32 ± 0.13 SE; Table S6). This correlation was significantly different from 0 (χ^2^_(1)_= 24.18, p<0.001) and 1 (χ^2^_(1)_= 73.17, p<0.001). A GWAS found four SNPs in a single region at the distal end of chromosome 18 (68,320,039 to 68,448,747 bp; SNPs *oar3_OAR18_68320039, oar3_OAR18_68398710, oar3_OAR18_68401733* and *oar3_OAR18_68448747*) were significantly associated with 25(OH)D_2_ concentrations (MAF = 0.219, χ^2^_(1)_ = 49.95, λ-corrected P = 5.33 × 10^−7^; Figures 2 and S5, Table S7). An animal model with genotype at *oar3_OAR18_68320039* fitt as an additional fixed factor found that genotypes AC and CC increased 25(OH)D_2_ concentrations by 0.91 (± 0.72 SE) and 2.40 (± 0.73 SE) nmol/l, respectively (Wald P = 2.12 × 10^−6^; note the AA and AC genotypes were not significantly different). This window contained three unannotated protein-coding regions, orthologous to the genes *TEX22, TEDC1* and *TMEM121*. None of these genes have previously been associated with vitamin D; similarly, there were no further candidate genes/orthologues within 1Mb of this association. Inclusion of a regional GRM in the animal model for 25(OH)D_2_ significantly improved the model (χ^2^_(1)_ = 10.31, p=0.001), showing that the chromosome 18 association region explained 6.56% (SE = 4.10%) of the phenotypic variance and 14.44% (SE = 4.06) of the additive genetic variance underlying 25(OH)D_2_ concentrations. No SNPs were associated with total 25(OH)D or 25(OH)D_3_ concentrations at the genome-wide level.

**Figure 2.**
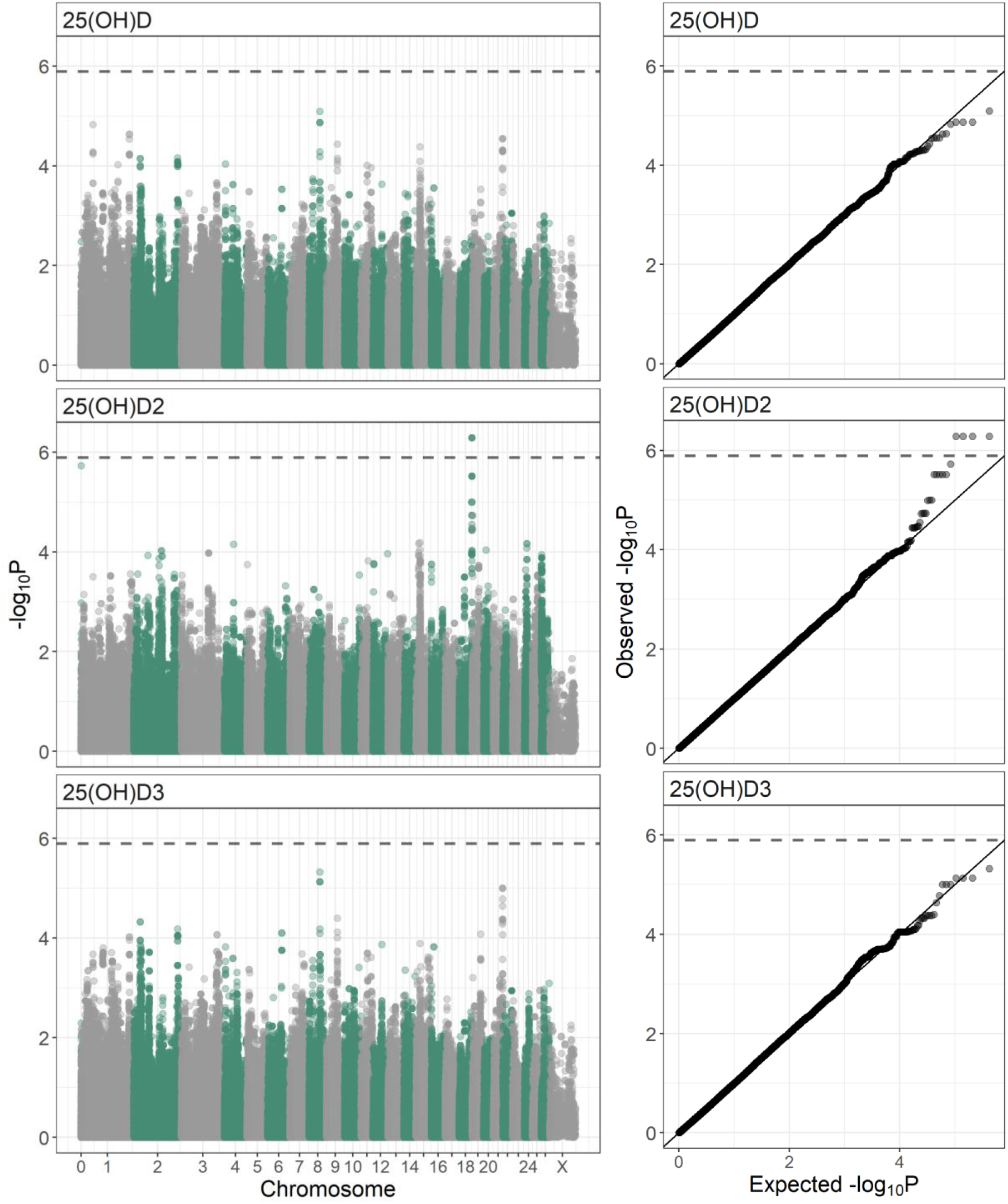
Genome-wide association results for 25(OH)D, 25(OH)D_2_, 25(OH)D_3_ concentrations at 420,556 SNPs in 900 Soay sheep across all ages. Left panel: Manhattan plots of p-values are corrected for inflation (by dividing by λ – 25(OH)D: 2.16, 25(OH)D_2_: 1.94, 25(OH)D_3_: 2.14) and the dashed line indicates the genome-wide significance threshold. Right panel: P-P plots showing the association between expected and observed (λ corrected) P-values where the solid line indicates the one to one line and the dashed line indicates the significance threshold.

### Associations between vitamin D concentrations and individual fitness

There was no evidence of an association between vitamin D measures in lambs and their subsequent first winter survival (Table 1). However, in adults, there was a significant interaction between 25(OH)D_2_ and sex on over-winter survival (χ^2^_(1)_ =4.790, p=0.029, Table 1 & Table S8, Figure 3), in which females with higher 25(OH)D_2_ concentrations in August were more likely to survive to the following year independently of their age and weight (females only model: b = 0.607 ± 0.182 SE, χ^2^_(1)_=12.119, p<0.001, Table S8). However, there was no significant association between 25(OH)D_2_ concentrations and winter survival for adult males (males only model: b = 0.177 ± 0.246, χ_(1)_^2^ =0.522, p=0.470, Table S8).

**Table 1.**
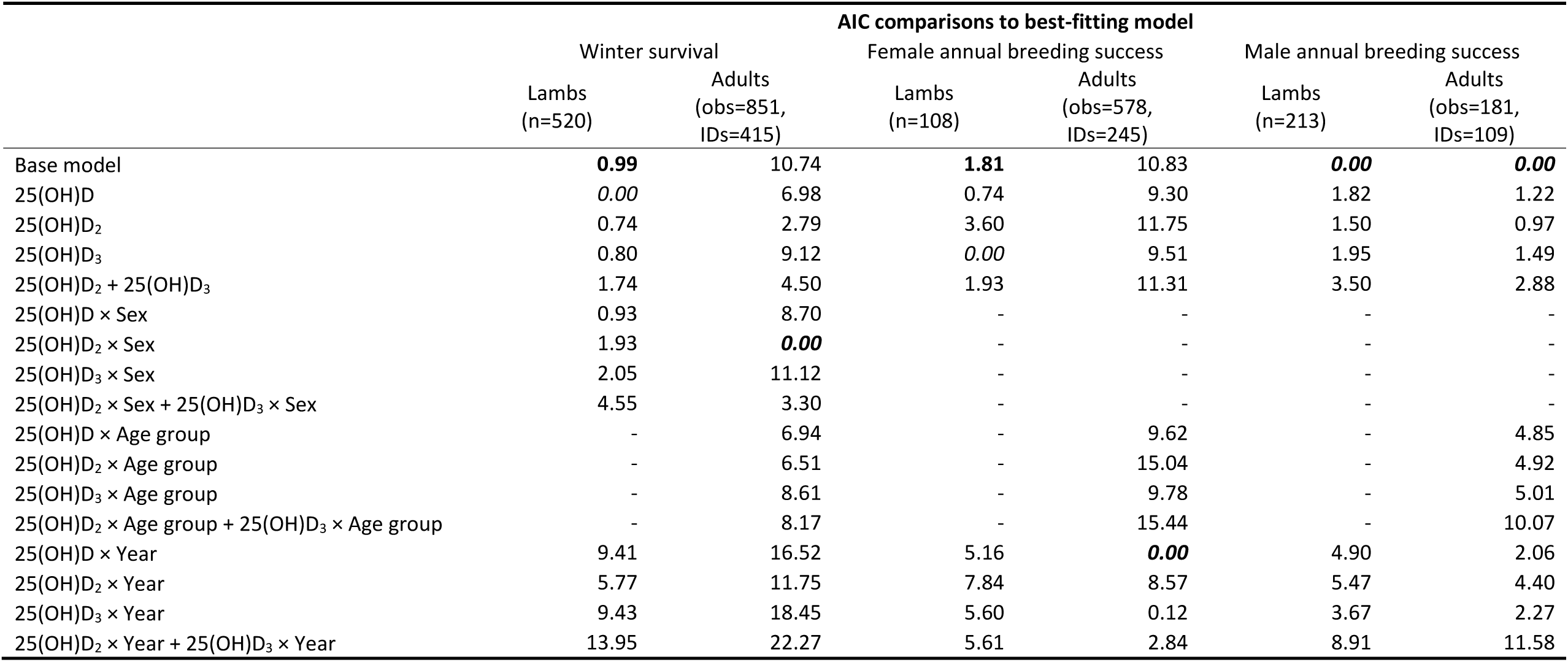
AIC comparison of over-winter survival and annual breeding success models in Soay sheep. The best-fitting model (where ΔAIC = 0) is highlighted in italics, and the most parsimonious model is highlighted in bold where ΔAIC <2 to the best-fitting model. All models with interactions also include these variables separately as main effects.

|  | AIC comparisons to best-fitting model |  |  |  |  |  |
| --- | --- | --- | --- | --- | --- | --- |
|  | Winter survival |  | Female annual breeding success |  | Male annual breeding success |  |
|  | Lambs<br>(n=520) | Adults<br>(obs=851,<br>IDs=415) | Lambs<br>(n=108) | Adults<br>(obs=578,<br>IDs=245) | Lambs<br>(n=213) | Adults<br>(obs=181,<br>IDs=109) |
| Base model | **0.99** | 10.74 | **1.81** | 10.83 | **0.00** | **0.00** |
| 25(OH)D | *0.00* | 6.98 | 0.74 | 9.30 | 1.82 | 1.22 |
| 25(OH)D<sub>2</sub> | 0.74 | 2.79 | 3.60 | 11.75 | 1.50 | 0.97 |
| 25(OH)D<sub>3</sub> | 0.80 | 9.12 | *0.00* | 9.51 | 1.95 | 1.49 |
| 25(OH)D<sub>2</sub> + 25(OH)D<sub>3</sub> | 1.74 | 4.50 | 1.93 | 11.31 | 3.50 | 2.88 |
| 25(OH)D × Sex | 0.93 | 8.70 | - | - | - | - |
| 25(OH)D<sub>2</sub> × Sex | 1.93 | **0.00** | - | - | - | - |
| 25(OH)D<sub>3</sub> × Sex | 2.05 | 11.12 | - | - | - | - |
| 25(OH)D<sub>2</sub> × Sex + 25(OH)D<sub>3</sub> × Sex | 4.55 | 3.30 | - | - | - | - |
| 25(OH)D × Age group | - | 6.94 | - | 9.62 | - | 4.85 |
| 25(OH)D<sub>2</sub> × Age group | - | 6.51 | - | 15.04 | - | 4.92 |
| 25(OH)D<sub>3</sub> × Age group | - | 8.61 | - | 9.78 | - | 5.01 |
| 25(OH)D<sub>2</sub> × Age group + 25(OH)D<sub>3</sub> × Age group | - | 8.17 | - | 15.44 | - | 10.07 |
| 25(OH)D × Year | 9.41 | 16.52 | 5.16 | **0.00** | 4.90 | 2.06 |
| 25(OH)D<sub>2</sub> × Year | 5.77 | 11.75 | 7.84 | 8.57 | 5.47 | 4.40 |
| 25(OH)D<sub>3</sub> × Year | 9.43 | 18.45 | 5.60 | 0.12 | 3.67 | 2.27 |
| 25(OH)D<sub>2</sub> × Year + 25(OH)D<sub>3</sub> × Year | 13.95 | 22.27 | 5.61 | 2.84 | 8.91 | 11.58 |

**Figure 3.**
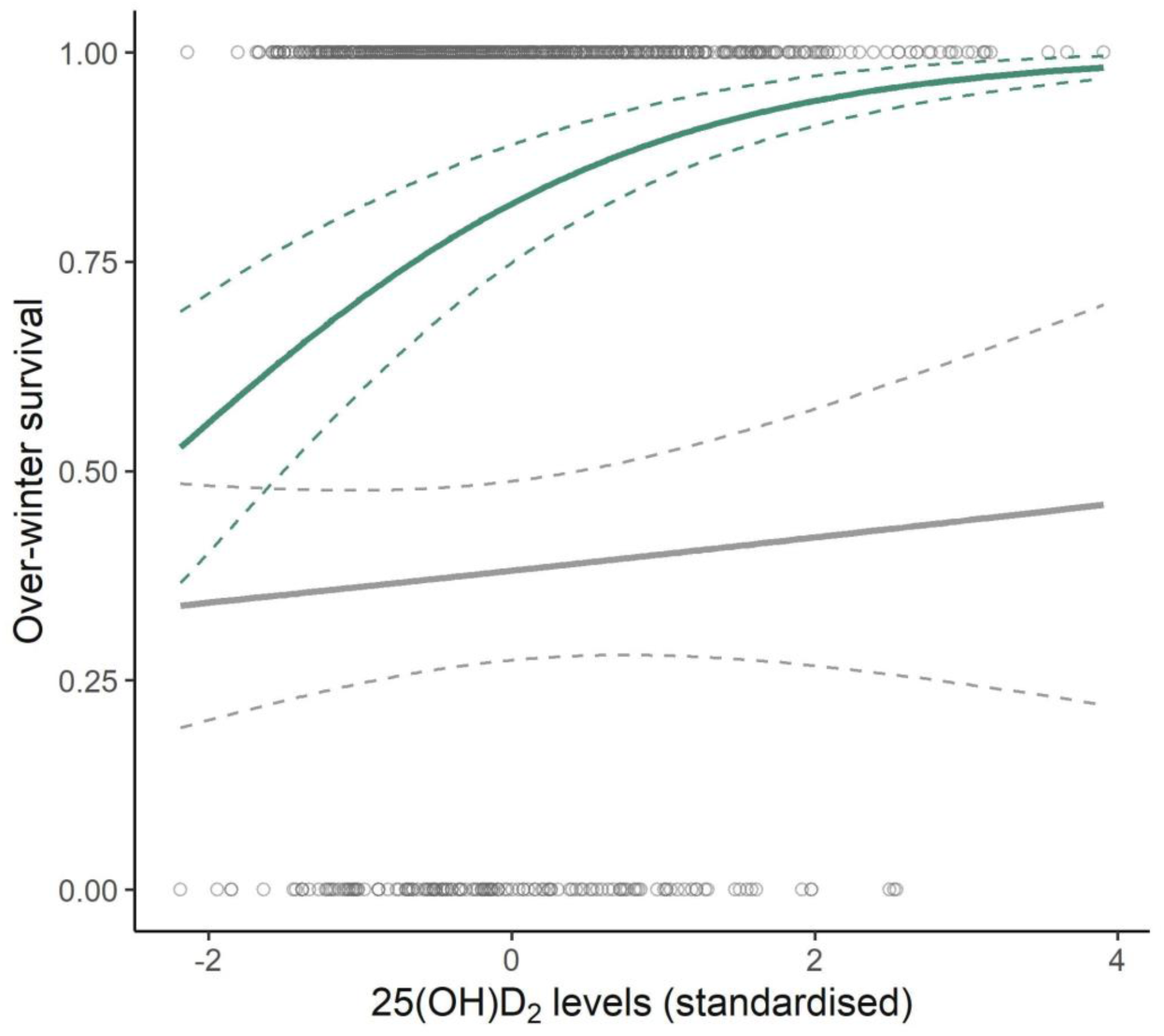
Scatterplot of raw data and general linear model predictions for associations between 25(OH)D_2_ concentrations (standardised, for full details see Methods) and subsequent over-winter survival for adult female (green) and male (grey) Soay sheep. The slope is predicted from the model with both sexes (Table S8) with other fixed effects set as follows: yearling age group, dark coat colour, the year 2011 and average weight. The dashed lines represent standard errors.

In females, there was no association between vitamin D measures of lambs and the probability of breeding in their first year (Table 1). However, in adult females, there was a significant interaction between 25(OH)D and 25(OH)D_3_ and year on annual breeding success (25(OH)D × Year: χ^2^_(5)_=19.304, p=0.002, 25(OH)D_3_ × Year: χ^2^_(5)_=19.389, p=0.002, Table 1 & Table S9, Figure 4). Total 25(OH)D and 25(OH)D_3_ concentrations in August positively predicted a female’s fecundity the following spring, but only in certain years (Table S9 & S10, Figure 4). In 2012, both total 25(OH)D and 25(OH)D_3_ concentrations positively predicted a female’s fecundity the following spring and in 2016 there was a marginally significant association between 25(OH)D_3_ concentrations and female fecundity (Table S10, Figure 4). For males, there was no evidence for associations between any of the vitamin D measures and the probability of males siring offspring in their first year, or annual breeding success in subsequent years for adult males (Table 1).

**Figure 4.**
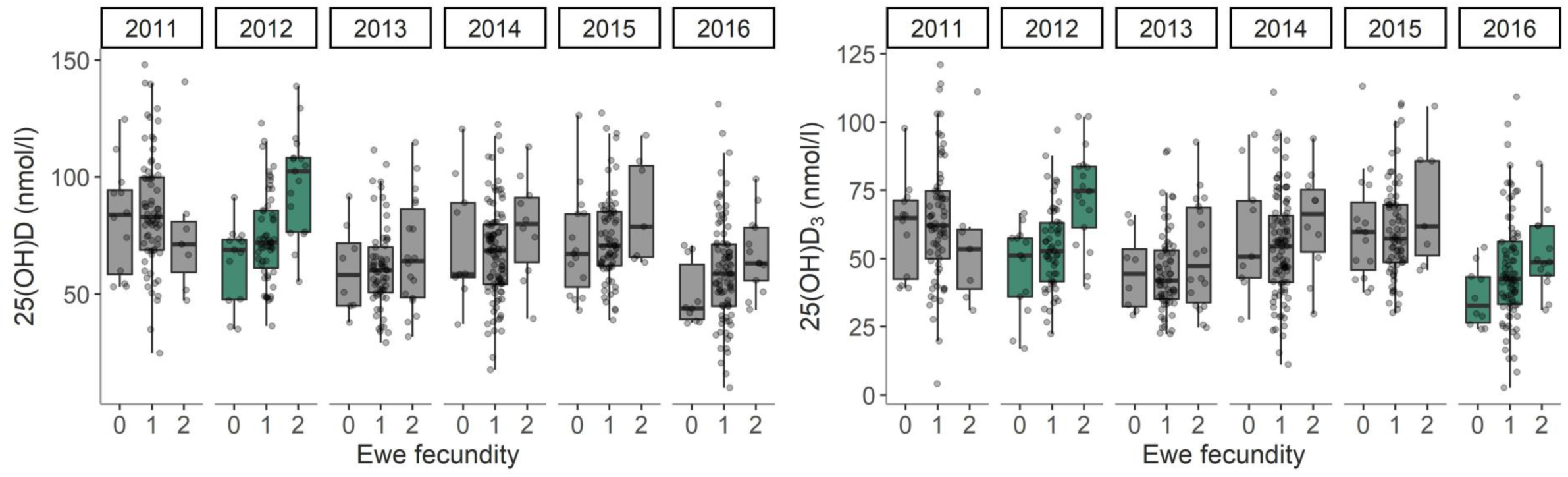
Boxplots of raw data showing associations between total 25(OH)D and 25(OH)D_3_ plasma concentrations and adult female fecundity by year in Soay sheep. Green boxplots indicate years in which there was a significant association (p<0.05, Table S10) between total 25(OH)D or 25(OH)D_3_ and female fecundity.

## Discussion

This study investigated the genetic architecture underlying vitamin D status in a wild mammal population and associations with fitness under natural conditions. We demonstrate that vitamin D status is repeatable within the lifetime of individuals and moderately heritable. We found limited evidence that heritable variation in vitamin D status was underpinned by loci of major phenotypic effect: a single genomic region explained ∼6.6% of the phenotypic variance in 25(OH)D_2_, but almost all heritable variation in circulating total 25(OH)D, 25(OH)D_2_ and 25(OH)D_3_ concentrations appears to be driven by many loci of small effect. Further, we have shown that vitamin D status is under age-, sex- and environment-dependent selection and selection patterns differed between cutaneous (25(OH)D_3_) and orally-derived sources (25(OH)D_2_) of vitamin D. Diet-derived vitamin D metabolite concentrations (25(OH)D_2_) were positively associated with female adult survival across all years whereas cutaneously-produced concentrations (25(OH)D_3_) predicted female fecundity in certain good environment years. Our study provides evidence that vitamin D status is heritable and has the potential to respond to selection under natural conditions and offers a unique insight into how vitamin D metabolism is associated with fitness in the wild.

We found that all vitamin D metabolites were relatively repeatable within individuals, despite the environmental heterogeneity experienced by individuals in this population. Longitudinal studies of vitamin D status in humans have similarly found that total 25(OH)D concentrations are correlated across subsequent sampling points within individuals ^37–41^. However, such studies are complicated by a more varied dietary intake in vitamin D and the greater ability of individuals to modify their UV exposure by modifying their behaviour and geographical location than in our wild mammal population. In our study, between 59-87% of the among-individual variation (repeatability) was due to additive genetic variation. Although vitamin D status is expected to be heavily influenced by environment, namely diet for 25(OH)D_2_ and sunlight for 25(OH)D_3_, our data show that there is still considerable genetic variation contributing to variation in circulating concentrations of total 25(OH)D, with heritabilities of 16.3% to 18.8%. Previous estimates from human populations using twin or familial studies have estimated the heritability of total 25(OH)D as high as 90% ^14,15,42–44^, but more recent SNP-based heritabilities have been modest, ranging from 7.5% to 13% ^11,13,45^. Our results provide evidence that total 25(OH)D as well as 25(OH)D_2_ and 25(OH)D_3_ are heritable in Soay sheep, indicating these traits have the potential to respond to selection and provide an important basis for further investigations into how genetic variation is maintained in these traits, including genotype-by-environment interactions.

The evolutionary potential of a given trait is also dependent on genetic correlations with other traits which may otherwise be masked at the phenotypic level ^46^. No previous studies have investigated the genetic correlation between diet-derived 25(OH)D_2_ and cutaneously produced 25(OH)D_3_ concentrations, and this cannot be readily interpreted in human populations due to the widespread consumption of vitamin D_3_ containing foodstuffs ^2^. Our study found a moderate positive genetic correlation between the two differentially obtained metabolites of 0.322, suggesting that variation in plasma concentrations of 25(OH)D_2_ and 25(OH)D_3_ are under some degree of shared genetic control. While selection could act independently on the two metabolites, this observed genetic correlation will constrain the evolution of one of the traits if antagonistic selection were to occur. Given the differential sources of these two metabolites, there are likely to be differences in the genetic pathways or networks involved in the metabolism of these two traits. For example, diet-derived 25(OH)D_2_ concentrations could be associated with genes underlying foraging behaviour and competitive ability and, cutaneously derived 25(OH)D_3_ may be related to differences in behaviour with respect to sheltering and exposure to sunlight. Future studies of individual variation in UV exposure and diet selection in unmanaged and unsupplemented populations like the Soay sheep could give more insight into the natural drivers of genetic variation in vitamin D metabolism.

A genome wide association study identified a single genomic region on chromosome 18 explaining ∼6.6% of the phenotypic variance and ∼14.% of the additive genetic variance in 25(OH)D_2_. Significant SNPs on chromosome 18 did not correspond directly to any compelling candidate genes associated with total 25(OH)D concentrations in humans ^9–13^. Two genes within the broader region, *PPP1R13B* and *DLK1* (∼1.28 Mb and ∼2.67 Mb away, respectively) were recently implicated in a human GWAS of circulating 25(OH)D levels ^13^. There was moderate LD between SNPs corresponding to *PPP1R13B* and the most highly associated SNPs (r^2^ = 0.765), suggesting that there may be some element of shared architecture of this trait. However, with the exception of this association, almost all heritable variation in vitamin D concentrations appears to be have a polygenic basis i.e. variation is driven by many genes of small effect throughout the genome. The high levels of LD in the Soay sheep genome aid the detection of trait loci using GWAS approaches. However our sample sizes are considerably lower than those of human studies ^13,16^, which may reduce the power to detect associations of moderate effect sizes and/or at alleles of low frequency within the population. Nevertheless, previous GWA studies of body size, antibody levels and recombination rate using only one tenth of the current SNP marker dataset successfully identified multiple loci explaining 2.6 to 46.7% of the phenotypic variance in these traits ^26,47,48^. Consequently, it is unlikely that there are major effect loci underpinning total 25(OH)D, 25(OH)D_2_ or 25(OH)D_3_ plasma concentrations in the Soay sheep. A previous study in this system documented that ewes with a light coat had higher total 25(OH)D and 25(OH)D_3_ concentrations than ewes with a dark coat colour ^18^ and this association was confirmed in both sexes in this six year dataset. The light/dark coat colour polymorphism is determined by a single base pair substitution at the *TYRP1* locus on chromosome 2 ^49^. This region was not associated with vitamin D status in the GWAS, suggesting that coat colour in itself does not explain enough variation in vitamin D status in this population for this region to have been detected.

In this study we were able to test for associations between 25(OH)D and mortality for the first time in a wild population, and were able to dissect associations between total (25(OH)D), oral (25(OH)D_2_) and cutaneously-derived (25(OH)D_3_) sources of vitamin D. Previous studies in both human and companion animals have demonstrated an association between low total 25(OH)D concentrations and poor health outcomes, including mortality ^2–4,50^. Interestingly, in this population we found a positive association between diet-derived 25(OH)D_2_ and overwinter survival in adult female sheep, but no association between total 25(OH)D or 25(OH)D_3_ concentrations during August and subsequent overwinter survival. This association between 25(OH)D_2_, but not total 25(OH)D or 25(OH)D_3_ summer concentrations, and female overwinter survival could be indicative of the environment experienced by the study population: no individuals receive any vitamin D_3_ supplemented food overwinter and individuals live at a latitude where cutaneous production of vitamin D_3_ is not possible between November to March which is the time period where mortality is highest on the island ^19,51,52^. Consequently, 25(OH)D_2_ may constitute almost all of the total winter total 25(OH)D concentration and so it is perhaps unsurprising that 25(OH)D_2_ rather than 25(OH)D_3_ or total 25(OH)D is linked to over-winter survival. Prolonged low concentrations of total 25(OH)D are likely to be highly deleterious to sheep with metabolic consequences including hypocalcaemia and skeletal disorders ^53^. This is particularly pertinent following the growing awareness that rickets remains a significant health issue in extensively farmed hill sheep farmed at a similar latitude to the Soay population ^54^. This case series highlighted that cutaneously derived 25(OH)D_3_ is often very low in late winter in Scottish hill sheep and so if insufficient vitamin D_2_ is consumed during the winter period then pronounced hypovitaminosis D with associated metabolic and skeletal complications can develop ^54^.

The lack of an association between summer vitamin D status and over winter survival in males may be due to the smaller sample size of males in the study or because males rut before the winter and forgo grazing ^19^. As a result, a summer measure of grazing ability may be unlikely to extend to the rut or reflect the status and condition of males going into the winter. Further, the absence of an association in lambs may be because they were just weaned at the time of blood sampling and were unlikely to be grazing as much as older animals as observed by their lower plasma concentrations of 25(OH)D_2_. While it is unclear whether 25(OH)D_2_ is acting as a marker of foraging capability or is directly leading to ill-health and contributing to death risk, our study provides evidence of an association between vitamin D status and subsequently mortality in a wild animal, and suggests that selection could be acting in different ways on 25(OH)D_2_ and 25(OH)D_3_ concentrations.

Higher plasma concentrations of vitamin D in August predicted increased fecundity in females, but not males, the following spring. This relationship was predominantly driven by 25(OH)D_3_ and was highly year dependent; the association was observed in 2012 and was marginally significant in 2016. Previously we identified a positive association between serum 25(OH)D concentrations and female fecundity using data from 2012 only ^18^. Our study recapitulates this finding using plasma samples, but also suggests that this finding is only apparent under certain ecological conditions. In 2012, when the relationship between 25(OH)D and 25(OH)D_3_ and ewe fecundity was highly significant, there was a low sheep density and low competition for resources during gestation and lactation following a population crash (Figure S6). This suggests there is environment-dependent fecundity selection for vitamin D status under benign ecological conditions. Our previous study determined that the relationship between vitamin D status and reproductive success is driven by increased fecundity rather than improved post parturition maternal care and lamb survival ^18^. The finding that increased vitamin D status is associated with improved female reproductive outcomes is consistent with findings in experimental models that vitamin D deficiency is linked to infertility, reduced pregnancy rates and litter sizes ^5,6,55–58^. Conversely, the lack of an association between vitamin D status and male breeding success is unsurprising given the limited evidence that vitamin D supplementation influences male reproductive outcomes in humans ^58–60^. Our results provide evidence for both sex and environment-dependent selection on vitamin D status, and suggests possible mechanisms by which genetic variation may have been maintained in these traits.

In conclusion, our study provides the first insights into the genetic basis and selection on vitamin D status in a wild population. Further study is warranted to investigate how well summer vitamin D concentrations predict winter vitamin D concentrations and whether selection patterns on winter vitamin D metabolites differ. Crucially, further studies in this population are needed to establish whether there are direct effects of vitamin D status on mortality and fecundity, or whether these measures are picking up signals of correlated traits such as diet choice, foraging capability or other aspects of behaviour that are themselves associated with fitness.

## Supporting information

Supplemental table S7 - GWAS Results for Vitamin D status.

## Acknowledgements

We thank Ian Stevenson and all Soay sheep project members and volunteers for collection of data and samples. Phil Ellis, Camillo Bérénos and Hannah Lemon prepared DNA samples for Ovine SNP50 BeadChip genotyping. Louise Evenden, Jude Gibson and Lee Murphy carried out SNP genotyping at the Wellcome Trust Clinical Research Facility Genetics Core, Edinburgh. The pedigree was constructed with advice from Jisca Huisman, and imputation was previously carried out by Martin Stoffel. This work has made use of the Edinburgh Compute and Data Facility (http://www.ecdf.ed.ac.uk/). Permission to work on St Kilda is granted by The National Trust for Scotland, and logistical support was provided by QinetiQ, Eurest and Kilda Cruises. This work was supported by grants from NERC, ERC, BBSRC and the Royal Society.

## Author Contributions

This study was conceived by AMS, RJM and DHN. JMP, JGP and DHN manage the long-term Soay sheep study system. Samples were collected by JMP and JGP. Lab work was undertaken by JB. AMS and SEJ performed all data analyses with inputs from DHN, IH and RJM. AMS wrote the first draft of the manuscript with input from SEJ, RJM and DHN. All authors provided comments on the manuscript and gave final approval for publication.

