## Supplemental Figures S1-S6 and Tables S1-S6 and S8-S10 for "Vitamin D status is heritable and under environment-dependent selection in the wild"

### Supplementary Figures and Tables

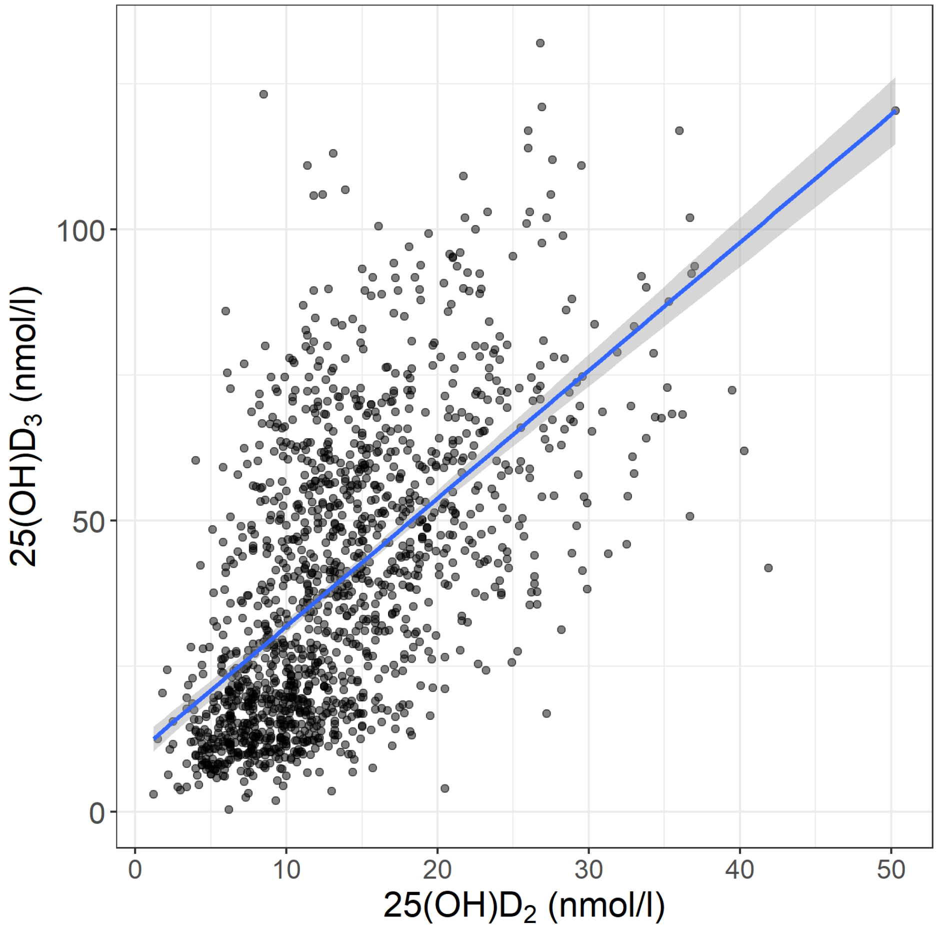


**Figure S1.** Scatterplot of the correlation between 25(OH)$D_2$ and 25(OH)$D_3$ plasma concentrations in Soay sheep.

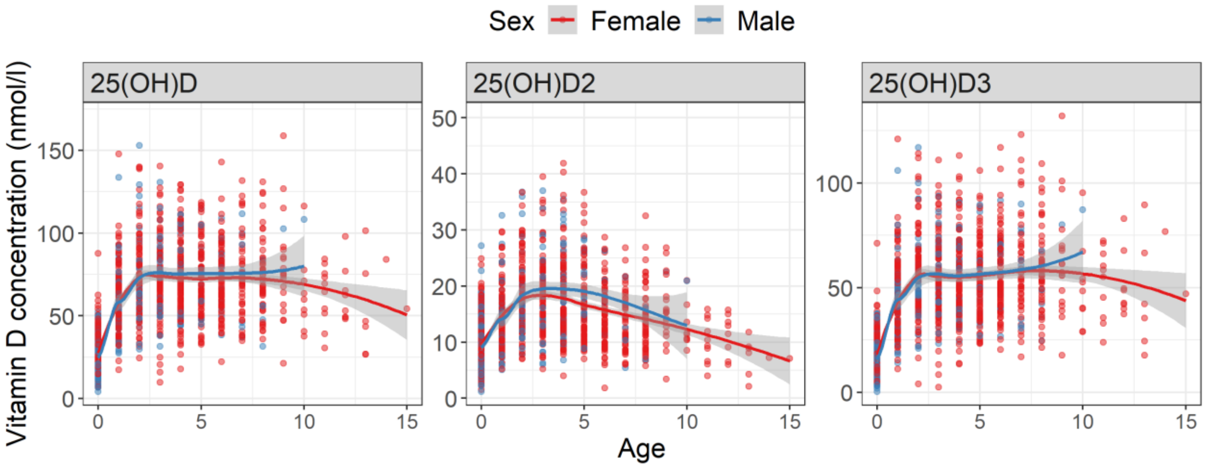


**Figure S2.** Plots of total 25(OH)D, 25(OH)$D_2$ and 25(OH)$D_3$ plasma concentrations with age using the raw data. Curves are shown for females (red) and males (blue) using a loess smoothing method in ggplot2 v3.3.0 [61] in R v3.6.2.

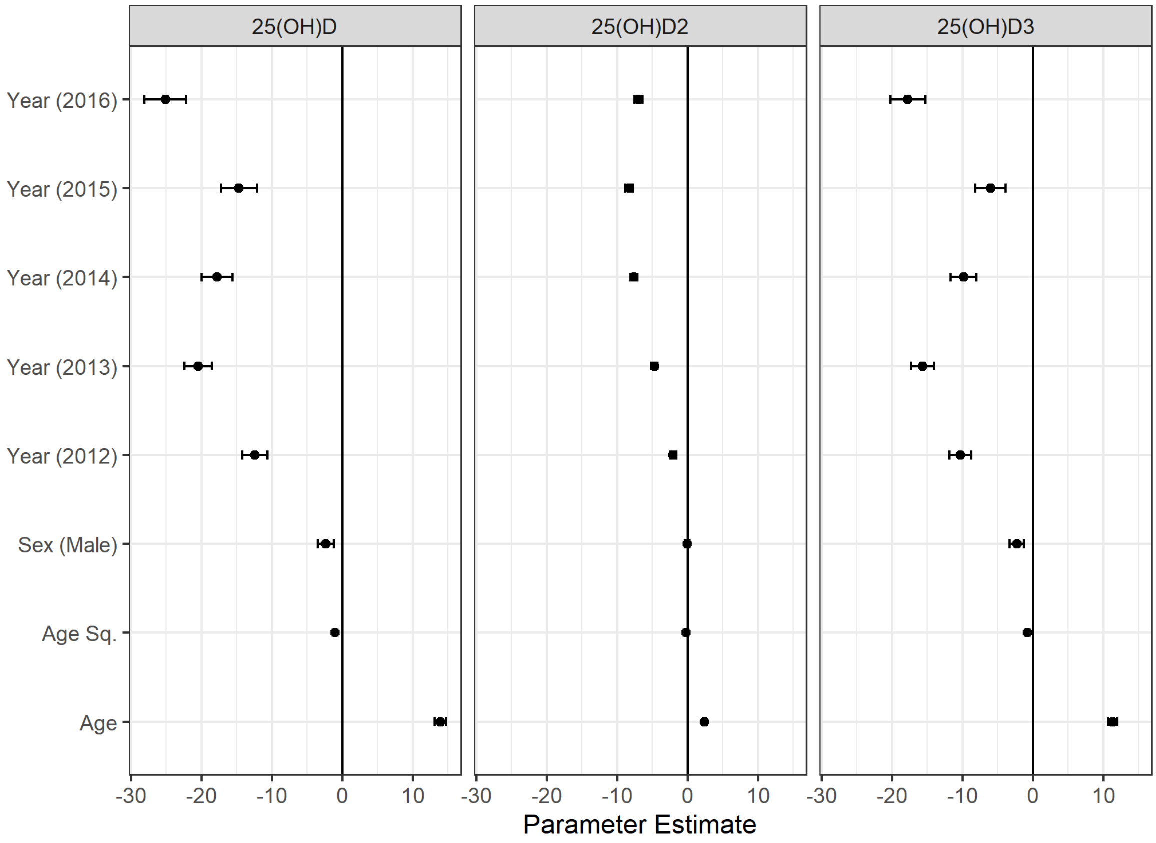


**Figure S3.** Parameter estimates and 95% confidence intervals of the fixed effects in the animal model (from Table S4) relative to the model intercept (set to 0). Reference levels are set as follows: sex (female) and year (2011).

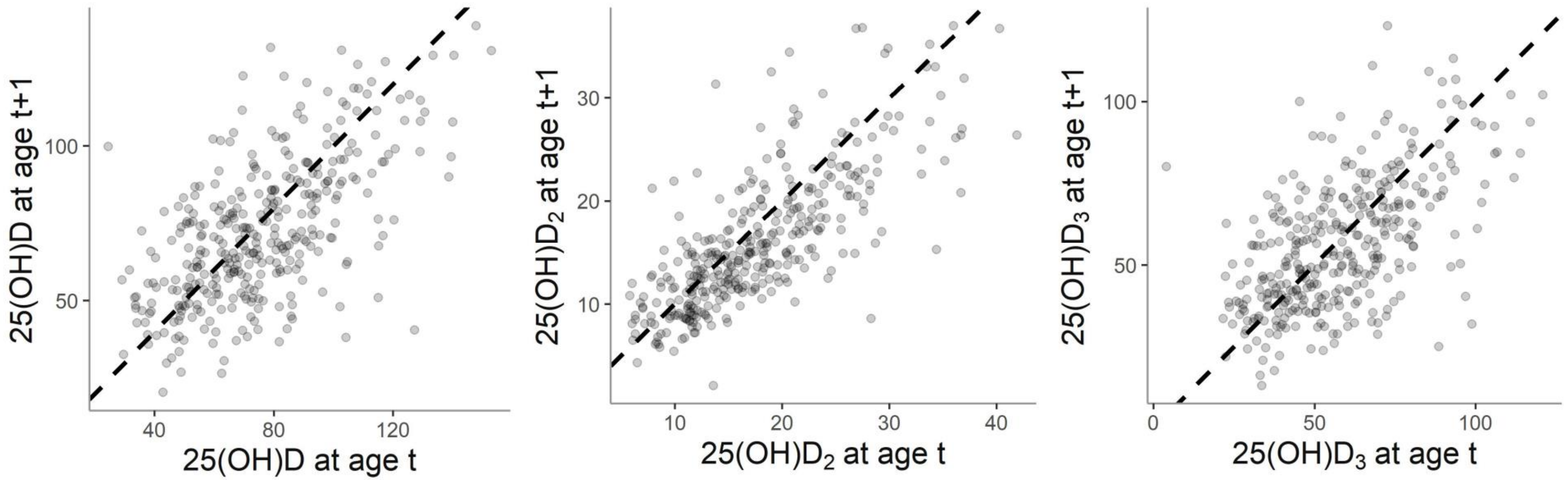


**Figure S4.** Temporal correlation in total 25(OH)D, $25(OH)D_2$ and $25(OH)D_3$ plasma concentrations (nmol/l) in adult Soay sheep. Scatterplots show raw data for adult sheep (≥1 year) which had two vitamin D measures in two consecutive years with a dashed line indicating a perfect 1:1 relationship.

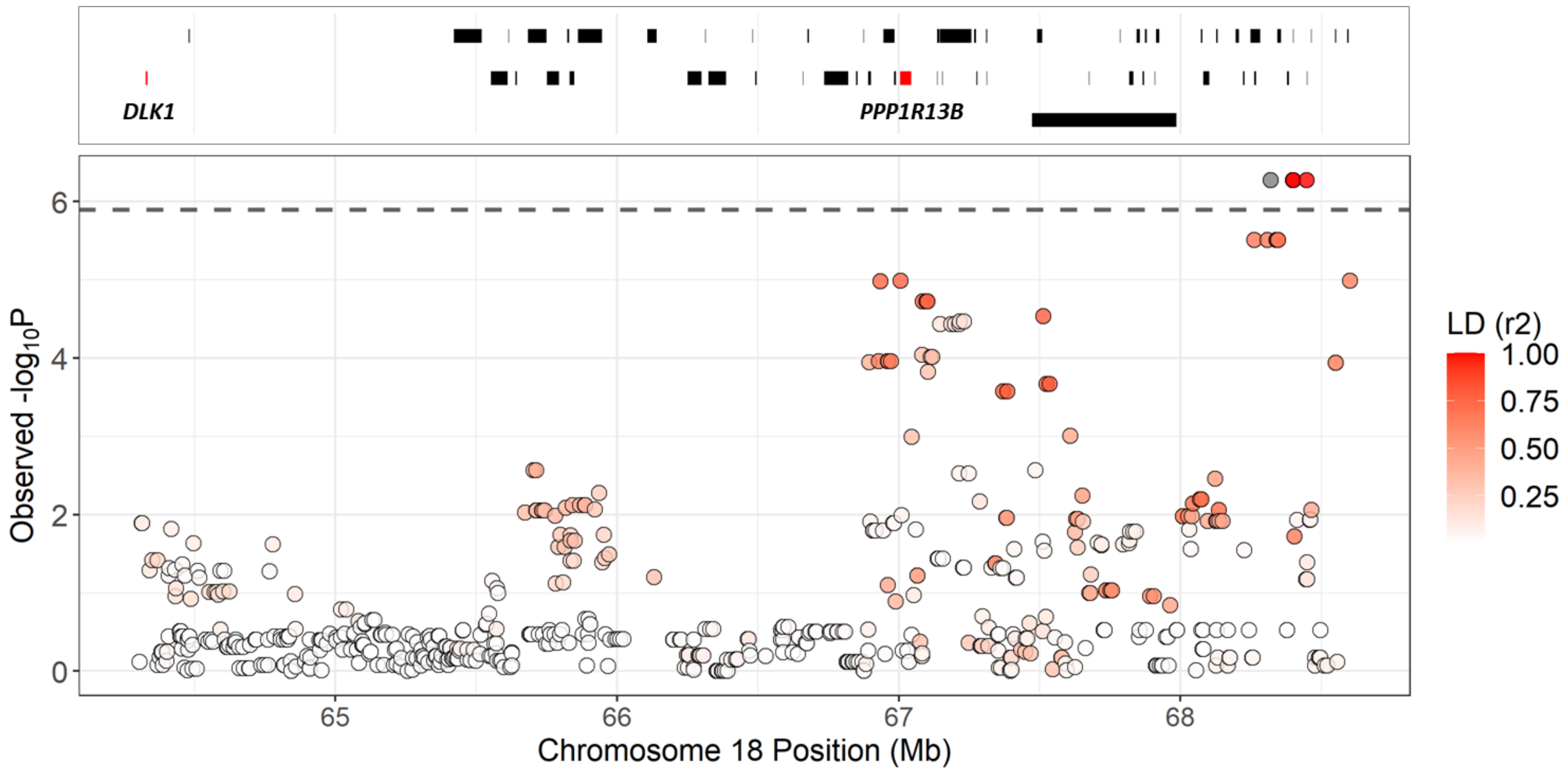


**Figure S5.** Local association of 25(OH)$D_2$ concentrations at the most highly associated region on chromosome 18. The dotted line indicates the genome-wide significance threshold equivalent to an experiment-wide threshold of P = 0.05. Points are colour-coded by their linkage disequilibrium with SNP ID *oar3_OAR18_68320039* indicated as a grey point ($r^2$ correlation, calculated using the *r2fast* function in GenABEL v1.8-0 [22]). Underlying data and effect sizes are provided in Table S7. Gene positions are shown in the grey panel at the top of the plot and were obtained from Ensembl (gene build ID Oar_v3.1.94). Genes coloured red have GO terms associated with vitamin D status or have previously been identified in human GWAS studies.

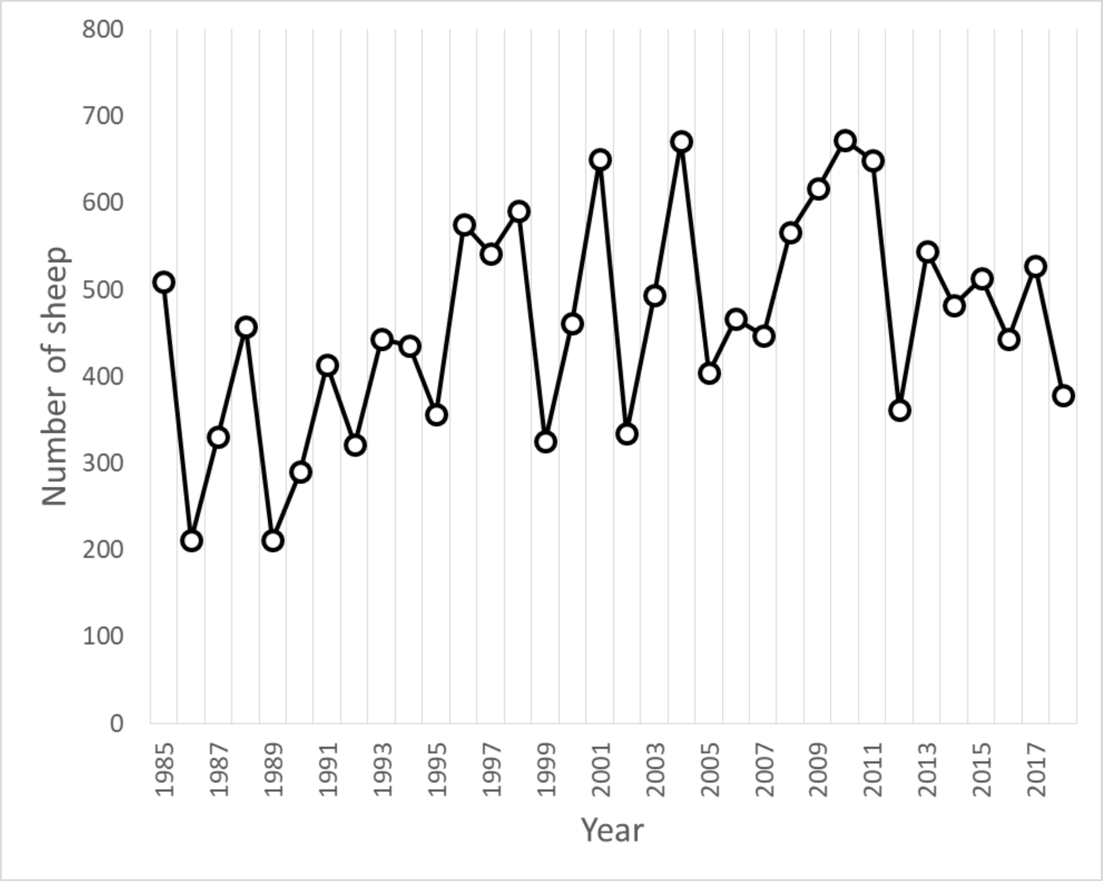


**Figure S6.** The number of Soay sheep in the study area in Village Bay on St Kilda by year.

**Table S1.** Study sample size breakdown by sex, age group and year for A) all samples and B) genotyped samples. Numbers in parentheses indicate the number of unique individuals.

**A. ALL SAMPLES**

| Sex | | | | | |
|---|---|---|---|---|---|
| Females | | | Males | | |
| 1000 (572) | | | 452 (345) | | |
| Age group | | | | | |
| Lambs | | Yearlings | 2-6 years | 7 years + | |
| 558 | | 152 | 555 (296) | 163 (102) | |
| Year | | | | | |
| 2011 | 2012 | 2013 | 2014 | 2015 | 2016 |
| 282 | 159 | 269 | 261 | 244 | 237 |

**B. GENOTYPED SAMPLES**

| Sex | | | | | |
|---|---|---|---|---|---|
| Females | | | Males | | |
| 962 (544) | | | 441 (336) | | |
| Age group | | | | | |
| Lambs | | Yearlings | 2-6 years | 7 years + | |
| 545 | | 147 | 548 (292) | 163 (102) | |
| Year | | | | | |
| 2011 | 2012 | 2013 | 2014 | 2015 | 2016 |
| 278 | 156 | 263 | 250 | 232 | 224 |

**Table S2.** AIC comparison of first year breeding success models for female and male lambs when data from the year 2011 was included in the models (see Methods). The best-fitting model (where ΔAIC = 0) is highlighted in italics, and the most parsimonious model is highlighted in bold where ΔAIC <2 to the best-fitting model. All models with interactions also include these variables separately as main effects.

| | **AIC comparisons to best-fitting model** | |
|---|---|---|
| | Annual breeding success | |
| | Females (n=112) | Males (n=260) |
| Base model | **1.81** | ***0.00*** |
| 25(OH)D | 0.74 | 1.82 |
| 25(OH)$D_2$ | 3.60 | 1.50 |
| 25(OH)$D_3$ | *0.00* | 1.95 |
| 25(OH)$D_2$ + 25(OH)$D_3$ | 1.93 | 3.50 |
| 25(OH)D × Year | 7.16 | 6.90 |
| 25(OH)$D_2$ × Year | 9.84 | 7.47 |
| 25(OH)$D_3$ × Year | 7.60 | 5.67 |
| 25(OH)$D_2$ × Year + 25(OH)$D_3$ × Year | 7.61 | 12.91 |

**Table S3.** Linear mixed model results of associations between sex, age, coat colour and year with total 25(OH)D, 25(OH)$D_2$ and 25(OH)$D_3$ concentrations in St Kilda Soay sheep. Included are the estimated effects (Estimate), standard error (SE) and the significance of fixed effects based on a likelihood ratio test (LRT, P-value) where all d.f.=1 except for year where d.f.=5. Sample size for all models is 1428 measures from 898 individuals.

| | **25(OH)D** | | | | **25(OH)$D_2$** | | | | **25(OH)$D_3$** | | | |
|---|---|---|---|---|---|---|---|---|---|---|---|---|
| variables | Estimate | SE | LRT | P-value | Estimate | SE | LRT | P-value | Estimate | SE | LRT | P-value |
| **fixed effects** | | | | | | | | | | | | |
| Intercept | 46.279 | 2.609 | - | - | 15.886 | 0.525 | - | - | 30.440 | 2.174 | - | - |
| Sex (male) | -2.142 | 1.225 | 3.123 | 0.077 | -0.186 | 0.319 | 0.359 | 0.549 | -1.995 | 1.047 | 3.699 | 0.054 |
| Age (years) | 13.758 | 0.784 | - | - | 2.355 | 0.173 | - | - | 11.290 | 0.659 | - | - |
| Age (years, quadratic) | -1.004 | 0.065 | 219.160 | <0.001 | -0.222 | 0.016 | 178.170 | <0.001 | -0.777 | 0.055 | 184.430 | <0.001 |
| Coat colour (light) | 5.512 | 1.354 | 16.526 | <0.001 | -0.742 | 0.353 | 4.419 | 0.036 | 6.253 | 1.158 | 28.829 | <0.001 |
| Year 2012 | -12.505 | 1.833 | 111.360 | <0.001 | -2.196 | 0.455 | 152.610 | <0.001 | -10.314 | 1.557 | 129.190 | <0.001 |
| Year 2013 | -20.445 | 1.890 | - | - | -4.792 | 0.453 | - | - | -15.627 | 1.600 | - | - |
| Year 2014 | -18.007 | 2.099 | - | - | -7.756 | 0.482 | - | - | -10.070 | 1.769 | - | - |
| Year 2015 | -15.149 | 2.426 | - | - | -8.333 | 0.537 | - | - | -6.631 | 2.039 | - | - |
| Year 2016 | -25.360 | 2.778 | - | - | -7.051 | 0.593 | - | - | -18.082 | 2.328 | - | - |
| **random effects** | | | | | | | | | | | | |
| ID | 113.529 | | | | 8.658 | | | | 84.761 | | | |
| Birth year | 56.356 | | | | 1.640 | | | | 38.252 | | | |
| Residual | 230.167 | | | | 14.306 | | | | 165.837 | | | |

**Table S4.** Animal model fixed effect estimates (Est and SE) and their associated significance levels (P) for 25(OH)D, 25(OH)$D_2$, 25(OH)$D_3$ concentrations in Soay sheep. Reference levels for the intercept are as follows: sex (female), and year (2011). All fixed effects are included as factors with age listed in years. The significance of fixed effects were determined using conditional Wald F-tests. Sample size for all models were 1,403 measures from 880 individuals.

| | Total 25(OH)D | | | 25(OH)$D_2$ | | | 25(OH)$D_3$ | | |
|---|---|---|---|---|---|---|---|---|---|
| | Est | SE | P | Est | SE | P | Est | SE | P |
| (Intercept) | 45.984 | 2.889 | | 15.242 | 0.533 | | 30.897 | 2.348 | |
| Age | 13.928 | 0.821 | < 0.001 | 2.411 | 0.174 | < 0.001 | 11.307 | 0.681 | < 0.001 |
| $Age^2$ | -1.017 | 0.065 | | -0.227 | 0.016 | | -0.779 | 0.055 | |
| Sex (Male) | -2.338 | 1.147 | < 0.001 | -0.078 | 0.305 | 0.9269 | -2.297 | 0.986 | < 0.001 |
| Year (2012) | -12.409 | 1.827 | < 0.001 | -2.075 | 0.454 | < 0.001 | -10.317 | 1.550 | < 0.001 |
| Year (2013) | -20.449 | 1.927 | | -4.729 | 0.454 | | -15.683 | 1.622 | |
| Year (2014) | -17.761 | 2.195 | | -7.645 | 0.486 | | -9.826 | 1.831 | |
| Year (2015) | -14.661 | 2.584 | | -8.309 | 0.543 | | -6.034 | 2.140 | |
| Year (2016) | -25.125 | 2.997 | | -6.976 | 0.602 | | -17.754 | 2.468 | |

**Table S5.** Animal model variance component estimates and their associated proportions for 25(OH)D, 25(OH)$D_2$, 25(OH)$D_3$ plasma concentrations in Soay sheep. Variances reported are the additive genetic variance ($V_A$), permanent environment variance ($V_{PE}$), maternal identity variance ($V_M$), birth year variance ($V_{BYEAR}$) and residual variance ($V_{\mathbf{R}}$). Included are the variance component estimates (Est) and the proportion of the total phenotypic variance explained by the term (Prop) with their associated standard errors in brackets. Mean and Var values are the mean and variance for the raw data, respectively. Sample size for all models were 1,403 measures from 880 individuals.

| Vitamin D Measure | Component | Est | Prop | P |
|---|---|---|---|---|
| **25(OH)D** | $V_A$ | 68.218 (15.908) | 0.163 (0.038) | < 0.001 |
| Mean = 53.047 | $V_{PE}$ | 9.857 (14.142) | 0.024 (0.034) | 0.429 |
| Var = 834.916 | $V_M$ | 38.207 (10.988) | 0.091 (0.027) | < 0.001 |
| | $V_{BYEAR}$ | 74.583 (36.49) | 0.178 (0.072) | < 0.001 |
| | $V_R$ | 227.046 (11.852) | 0.543 (0.054) | |
| **25(OH)$D_2$** | $V_A$ | 4.205 (1.088) | 0.172 (0.042) | < 0.001 |
| Mean = 13.509 | $V_{PE}$ | 2.87 (1.066) | 0.117 (0.044) | 0.001 |
| Var = 43.518 | $V_M$ | 1.423 (0.708) | 0.058 (0.029) | 0.026 |
| | $V_{BYEAR}$ | 1.724 (0.948) | 0.071 (0.036) | < 0.001 |
| | $V_R$ | 14.228 (0.777) | 0.582 (0.038) | |
| **25(OH)$D_3$** | $V_A$ | 56.649 (12.145) | 0.188 (0.04) | < 0.001 |
| Mean = 39.540 | $V_{PE}$ | 26.569 (8.015) | 0.088 (0.027) | 0.403 |
| Var = 600.796 | $V_M$ | 7.676 (10.461) | 0.025 (0.035) | < 0.001 |
| | $V_{BYEAR}$ | 47.333 (23.177) | 0.157 (0.065) | < 0.001 |
| | $V_R$ | 163.3 (8.538) | 0.542 (0.049) | |

**Table S6.** Animal model variance/covariance matrices for a bivariate model of 25(OH)$D_2$ and 25(OH)$D_3$ concentrations in Soay sheep. Covariances and correlations were estimated at the additive genetic, permanent environmental and residual level. Variances are on the diagonal, covariances are highlighted in italics on the lower off-diagonal and correlations are highlighted in bold on the upper off-diagonal with standard errors indicated in brackets.

| Additive genetic | | |
|---|---|---|
| | 25(OH)$D_2$ | 25(OH)$D_3$ |
| 25(OH)$D_2$ | 4.952 (1.105) | **0.322 (0.132)** |
| 25(OH)$D_3$ | *5.630 (2.818)* | 61.717 (12.504) |
| **Permanent environment** | | |
| | 25(OH)$D_2$ | 25(OH)$D_3$ |
| 25(OH)$D_2$ | 4.304 (1.053) | **0.392 (0.159)** |
| 25(OH)$D_3$ | *4.967 (2.666)* | 37.219 (11.234) |
| **Residual** | | |
| | 25(OH)$D_2$ | 25(OH)$D_3$ |
| 25(OH)$D_2$ | 14.491 (0.792) | **0.526 (0.028)** |
| 25(OH)$D_3$ | *26.310 (2.137)* | 172.789 (9.170) |

**Table S7.** Full genome-wide association results for total 25(OH)D, 25(OH)$D_2$, 25(OH)$D_3$ plasma concentrations in Soay sheep. A1 and A2 are the reference and alternate allele at each SNP. effB is the slope of the effect of allele A2, with the standard error se_effB. Chi2.1df and P1df is the association chi-squared statistic and associated P-value, respectively, before correction with genomic control λ. Pc1df is the corrected P-value after genomic control. Exp is the corresponding P-value for that SNP locus assuming a null distribution of P-values (see Figure 2). Q.2 is the minor allele frequency.

**[file - 2_Table_S7_GWAS_Results.zip]**

**Table S8.** Generalised linear model results of the best-fitting model for over-winter survival in adult Soay sheep for both sexes and males and females separately. Included are the estimated effects (Estimate), standard errors (SE) and significance of fixed effects based on a likelihood ratio test (LRT, df, P).

| | **Over-winter survival** | | | | | | | | | | | | | | |
|---|---|---|---|---|---|---|---|---|---|---|---|---|---|---|---|
| | All adults Obs=851, IDs =415 | | | | | Females only Obs=674, IDs =309 | | | | | Males only Obs=177, IDs =106 | | | | |
| Fixed effects | Estimate | SE | LRT | df | P | Estimate | SE | LRT | df | P | Estimate | SE | LRT | df | P |
| Intercept | 1.512 | 0.476 | - | - | - | 1.982 | 0.594 | - | - | - | -0.600 | 0.631 | - | - | [illegible] |
| Sex (male) | -1.997 | 0.502 | - | - | - | | | | | | | | | | |
| Age group (2-6) | -0.316 | 0.407 | - | - | - | -0.619 | 0.464 | 32.376 | 2 | <0.001 | -0.503 | 0.876 | 16.022 | 2 | [illegible] |
| Age group (7+) | -2.022 | 0.485 | - | - | - | -2.409 | 0.577 | - | - | - | -3.667 | 1.294 | - | - | [illegible] |
| Coat colour (light) | 0.081 | 0.251 | 0.103 | 1 | 0.748 | 0.100 | 0.286 | 0.123 | 1.000 | 0.726 | 0.110 | 0.565 | 0.038 | 1 | [illegible] |
| Weight | 0.882 | 0.214 | 18.244 | 1 | <0.001 | 1.192 | 0.304 | 16.855 | 1 | <0.001 | 0.656 | 0.311 | 4.684 | 1 | [illegible] |
| Year (2012) | 4.172 | 1.034 | 124.106 | 5 | <0.001 | 3.662 | 1.042 | 80.830 | 5 | <0.001 | 19.752 | 2458.658 | 48.644 | 5 | [illegible] |
| Year (2013) | 0.793 | 0.326 | - | - | - | 0.722 | 0.387 | - | - | - | 0.729 | 0.622 | - | - | [illegible] |
| Year (2014) | 2.401 | 0.391 | - | - | - | 2.644 | 0.489 | - | - | - | 1.886 | 0.703 | - | - | [illegible] |
| Year (2015) | 1.994 | 0.362 | - | - | - | 1.775 | 0.404 | - | - | - | 2.953 | 0.872 | - | - | [illegible] |
| Year (2016) | 4.042 | 0.651 | - | - | - | 3.589 | 0.673 | - | - | - | 20.089 | 1768.911 | - | - | [illegible] |
| 25(OH)D2 | 0.640 | 0.174 | - | - | - | 0.607 | 0.182 | 12.119 | 1 | <0.001 | 0.177 | 0.246 | 0.522 | 1 | [illegible] |
| Sex (male) x Age group (2-6) | -0.494 | 0.637 | 6.826 | 2 | 0.033 | | | | | | | | | | |
| Sex (male) x Age group (7+) | -2.277 | 0.919 | - | - | - | | | | | | | | | | |
| Sex (male) x 25(OH)D2 | -0.557 | 0.257 | 4.790 | 1 | 0.029 | | | | | | | | | | |

**Table S9**. Generalised linear mixed model results of the two best-fitting models for adult female fecundity in Soay sheep. Included are the estimated effects (Estimate), standard errors (SE) and significance of fixed effects based on a likelihood ratio test (LRT, df, P).

| | **Ewe fecundity** | | | | | | | | | |
|---|---|---|---|---|---|---|---|---|---|---|
| | 25(OH)D Estimate | SE | LRT | df | P | $25(OH)D_3$ Estimate | SE | LRT | df | P |
| Age group (2-6 years) | 0.711 | 0.607 | 7.881 | 2 | 0.019 | 0.680 | 0.607 | 8.532 | 2 | 0.014 |
| Age group (7+ years) | -0.421 | 0.770 | - | - | - | -0.519 | 0.775 | - | - | - |
| Coat colour (light) | -0.165 | 0.489 | 0.114 | 1 | 0.735 | -0.228 | 0.492 | 0.215 | 1 | 0.643 |
| Weight | 0.461 | 0.251 | 3.527 | 1 | 0.060 | 0.482 | 0.251 | 3.851 | 1 | 0.050 |
| Year (2012) | -0.106 | 0.533 | - | - | - | 0.254 | 0.504 | - | - | - |
| Year (2013) | 0.921 | 0.561 | - | - | - | 1.026 | 0.543 | - | - | - |
| Year (2014) | 0.338 | 0.519 | - | - | - | 0.395 | 0.497 | - | - | - |
| Year (2015) | -0.490 | 0.561 | - | - | - | -0.403 | 0.549 | - | - | - |
| Year (2016) | 0.585 | 0.587 | - | - | - | 0.647 | 0.558 | - | - | - |
| 25(OH)D | -0.333 | 0.349 | - | - | - | | | | | |
| 2012 x 25(OH)D | 1.982 | 0.509 | 19.304 | 5 | 0.002 | | | | | |
| 2013 x 25(OH)D | 0.692 | 0.506 | - | - | - | | | | | |
| 2014 x 25(OH)D | 0.338 | 0.468 | - | - | - | | | | | |
| 2015 x 25(OH)D | 0.628 | 0.510 | - | - | - | | | | | |
| 2016 x 25(OH)D | 0.802 | 0.475 | - | - | - | | | | | |
| $25(OH)D_3$ | | | | | | -0.273 | 0.348 | - | - | - |
| 2012 x $25(OH)D_3$ | | | | | | 2.039 | 0.530 | 19.389 | 5 | 0.002 |
| 2013 x $25(OH)D_3$ | | | | | | 0.742 | 0.522 | - | - | - |
| 2014 x $25(OH)D_3$ | | | | | | 0.322 | 0.461 | - | - | - |
| 2015 x $25(OH)D_3$ | | | | | | 0.399 | 0.491 | - | - | - |
| 2016 x $25(OH)D_3$ | | | | | | 0.793 | 0.470 | - | - | - |

**Table S10.** Generalised linear mixed model results of associations between 25(OH)D and 25(OH)$D_3$ concentrations and adult female fecundity when models were run on each year separately. Included are the estimated effects (Estimate), standard errors (SE) and significance of associations between 25(OH)D or 25(OH)$D_3$ and adult female fecundity based on a likelihood ratio test (LRT, df, P) from a proportional odds logistic regression model in the MASS package v7.3-51.1 (Venables and Ripley, 2002) including age group (factor), coat colour (factor) and weight as fixed effects.

| | Ewe fecundity | | | | | | | | | |
|---|---|---|---|---|---|---|---|---|---|---|
| | 25(OH)D | | | | | 25(OH)$D_3$ | | | | |
| Year/Model | Estimate | SE | LRT | df | P | Estimate | SE | LRT | df | P |
| 2011 | -0.146 | 0.261 | 0.314 | 1 | 0.575 | -0.132 | 0.259 | 0.260 | 1 | 0.610 |
| 2012 | 1.159 | 0.296 | 18.258 | 1 | <0.001 | 1.236 | 0.317 | 18.160 | 1 | <0.001 |
| 2013 | 0.202 | 0.287 | 0.494 | 1 | 0.482 | 0.325 | 0.300 | 1.182 | 1 | 0.277 |
| 2014 | 0.190 | 0.295 | 0.416 | 1 | 0.519 | 0.184 | 0.281 | 0.430 | 1 | 0.512 |
| 2015 | 0.450 | 0.307 | 2.175 | 1 | 0.140 | 0.249 | 0.287 | 0.756 | 1 | 0.385 |
| 2016 | 0.499 | 0.266 | 3.601 | 1 | 0.058 | 0.512 | 0.261 | 3.917 | 1 | 0.048 |
